# Age-Dependent Increase and Dysregulation of the Splicing Machinery in Acute Myeloid Leukemia of Older Patients

**DOI:** 10.1101/2025.07.04.663197

**Authors:** Jordan B. Burton, Kristin Pladna, Joanna Bons, Mark A. Watson, Christina D. King, Timothy S. Pardee, Birgit Schilling

## Abstract

Acute myeloid leukemia (AML) is characterized by age-related resistance to therapy and poor outcomes. Initial remission rates vary from 30-80% with current therapies, but relapses are common. The 5-year survival is 30% overall but less than 10% for those 60 years of age or older. Resistance to current therapies is the central clinical challenge. This study investigated how age influences the AML proteome in patients with increasing age. We analyzed bone marrow or leukapheresis samples from 14 AML patients, including 9 older (≥ 66 years) and 5 younger (≤ 58 years) individuals, using proteomic technologies (Data-Independent Acquisitions, DIA-MS). We identified 4,471 protein groups, of which 889 exhibited significant changes and regulation between the two age groups. Proteins upregulated in older patients were predominantly associated with DNA transcription and translation, while downregulated proteins were linked to metabolic, catabolic, and extracellular matrix or structural processes. Significant changes in older samples affected the splicing machinery. Significant alterations in several of these factors, suggests either dysregulated or elevated splicing activity and reduced transcription initiation factor activity in older AML patients. Our study provides novel insights into molecular changes during AML arising in the older adult, highlighting the importance of age-specific therapeutic strategies.

**Statement of Significance of the Study:** This study is significant as it investigates a devastating and highly aggressive cancer, acute myeloid leukemia (AML), which often presents in patients with very low 5-year survival rates (32%), however which is even lower in patients over 60 years old (10%). This study really focuses on the complications and severe outcome in older patients which is not well understood. This is particularly relevant as older AML patients often do not respond to the available treatments. Understanding the pathways and protein profiles that are dysregulated in older AML patients compared to younger AML patients may in the future help to develop more customized therapeutic interventions or allow for some precision treatments. AML patients were stratified into cohort groups by age (older vs younger) and the proteomic profiling highlighted the particularly strong dysregulation of the splicing machinery in the older AML patients. While alternative splicing is often observed in cancers, here the emphasis is pointed to the fact that the dysregulation of the spliceosome and other pathways are greatly exacerbated in the older AML patients, likely leading to even worse outcomes and poor response to treatments in the older patients.

## 1 | INTRODUCTION

Acute myeloid leukemia (AML) is an aggressive malignancy of the bone marrow where neoplastic myeloid precursors proliferate uncontrollably and suppress normal marrow function leading to marrow failure and death (1). Approximately 20,000 US citizens are diagnosed with AML every year. The 5-year survival rate is 32%, but only 10% in patients over the age of 60. Age is an important prognostic marker with older patients uniformly experiencing worse outcome. Even matched by molecular and cytogenetic risk group, older patients have lower remission rates and shorter survival when compared to younger patients (2). While AML can be diagnosed at any age, it is primarily an age-associated malignancy with a median age at onset of 66-71 years old (3). Additionally, risk factors for accelerated aging such as chemotherapy or radiation exposure increase the risk of acquiring AML.

AML arises from the hematopoietic stem cells (HSC) or early myeloid progenitors. HSCs must survive the entire lifespan to generate the approximately 200 billion blood cells needed daily. These long-lived cells undergo biochemical changes with aging, including the biological changes referred to as hallmarks of aging (4, 5), many of which are also closely linked to the progression and relation between aging and cancer (6). However, how AML response and clinical behavior are affected by the biochemical changes during aging is largely unknown. Studies have shown that there are differences in AML gene expression profiles by age at onset and that pediatric AML appears molecularly distinct from adult AML (7). Our lab has previously demonstrated that HSCs from older AML patients show impaired mitochondrial capacity and may benefit from mitochondrial targeted agents (8). In what other respects the biochemical hallmarks of aging influence AML arising in an older host is largely unknown. To explore what other biological differences may exist between older and younger AML patients we assessed the proteomes of primary patient samples using mass spectrometric data-independent acquisitions (DIA)(9).

Previous studies have investigated predominantly RNA expression or DNA structural variants in AML (10, 11), our study is specifically designed to uncover effects of aging on the proteome of AML cells from older and younger patients.

In this study, we compared proteomic abundances across old and young AML patient bone marrow aspirates or leukapheresis samples to identify protein signatures specific to aging in older AML patients. We demonstrated a comprehensive and quantitative proteomic data-independent acquisition workflow to gain novel biological insights into the contribution of age on AML. We identified enriched factors associated with eukaryotic translation initiation in younger AML patients and dysregulation in the splicing machinery in older AML patients.

## 2 | MATERIALS AND METHODS

### 2.1 | Chemicals

HPLC grade acetonitrile and water were procured from Burdick & Jackson (Muskegon, MI). Sigma-Aldrich (St. Louis, MO) supplied the methanol, urea, triethylammonium bicarbonate buffer (TEAB), dithiothreitol (DTT), iodoacetamide (IAA), phosphoric acid, formic acid, and sodium dodecyl sulfate (SDS). Protease/phosphatase inhibitor cocktail (PIC) was obtained from Thermo Fisher Scientific (Waltham, MA). The hydrophilic– lipophilic balance (HLB) 10 mg Oasis solid phase extraction cartridges used for desalting were purchased from Waters (Milford, MA), while the sequencing grade trypsin utilized for protein digestion was acquired from Promega (Madison, WI). Indexed Retention Time (iRT) peptides were sourced from Biognosys (Schlieren, Switzerland).

### 2.2 | Collection of human samples

Diagnostic samples from a total of 14 AML patients were analyzed. All patients signed written informed consent document prior to the sample being obtained under an Atrium Health Wake Forest Baptist IRB approved protocol (IRB#00011131). Of these, 5 samples were from patients younger than 60 years of age (Median 54, range 26-58) and were considered the young cohort. The remaining 9 samples were from patients older than 60 years of age (median 74.5, range 66-85) and were considered the older cohort. A total of three samples were obtained from leukapheresis procedures (1 in the older cohort and 2 in the younger). The median blast cell percentage was 75% in the younger cohort and 87% in the older cohort. In the older cohort 5/9 patients had a normal karyotype and while only one in the younger cohort did. Three patients in the older cohort had an adverse karyoptype by ELN 2022 criteria compared with two in the younger cohort. Unfortunately, mutational data was not available for the majority of the patient samples.

### 2.3 | Bone marrow aspiration or leukapheresis sample acquisition and processing

All marrow aspirates or leukapheresis samples were obtained during clinically indicated bone marrow biopsies or leukapheresis procedures following signed informed consent under a protocol approved by the Comprehensive Cancer Center of Atrium Health Wake Forest Baptist (IRB#00011131). Samples consisted of 3-5 mL of marrow aspirate or leukapheresis sample placed in an EDTA tube. Samples were then spun at ∼1500 rpm at RT for 10 minutes to pellet cells. Washed cells were resuspended in media and mononuclear cells isolated by Ficoll gradient. Following isolation cells were washed in media and resuspended in freeze media (10% DMSO in FBS) and placed at -80 °C. Upon use cells were rapidly thawed, spun at 800 rpm for 10 minutes at 4 °C degrees and washed with ice cold PBS x2. Cell pellets were then snap frozen and placed at -80°C until analysis.

### 2.4 | Protein digestion and desalting

Cell pellets solubilized in 100 µL 0.5% SDS in 100 mM TEAB with 1X PIC. BCA protein assay was performed with 1:3 dilution. Aliquots containing 100 µg were reduced with DTT, alkylated with IAA, precipitated, digested with trypsin in a 1:25 ratio for 1 h at 47°C, an additional aliquot of trypsin was added in the same ratio, allowing the proteins to digest overnight at 37 °C using the S-trap mini (Protifi) protocol. Samples were desalted with 10 mg Oasis HLB Cartridges and dried by centrifugal evaporation. Samples were resuspended in 100 µL 0.2% formic acid in water at a final concentration of 1 µg/µL. Each sample was spiked with indexed Retention Time Standards (iRT, Biognosys) following the manufacturer’s guidelines.(12)

### 2.5 | Mass spectrometric analysis

Reverse-phase HPLC-MS/MS data was acquired using a Waters M-Class HPLC (Waters, Massachusetts, MA) connected to a ZenoTOF 7600 (SCIEX, Redwood City, CA) with an OptiFlow Turbo V Ion Source (SCIEX) equipped with a microelectrode. The chromatographic solvent system consisted of 0.1% formic acid in water (solvent A) and 99.9% acetonitrile, 0.1% formic acid in water (solvent B). Digested peptides (1 µg) were loaded on onto a Luna Micro C18 trap column (20 x 0.30 mm, 5 µm particle size; Phenomenex, Torrance, CA) over a period of 2 minutes at a flow rate of 10 µL/min using 100% solvent A. Peptides were eluted onto a Kinetex XB-C18 analytical column (150 x 0.30 mm, 2.6 µm particle size; Phenomenex) at a flow rate of 5 µL/min using a 120 minute microflow gradient (as described below), with each gradient ranging from 5 to 32% solvent B. The following MS parameters were used for all acquisitions: ion source gas 1 at 10 psi, ion source gas 2 at 25 psi, curtain gas at 30 psi, CAD gas at 7 psi, source temperature at 200 °C, column temperature at 30 °C, polarity set to positive, and spray voltage at 5000 V. All human samples were acquired in data-independent acquisition mode (DIA), which is comprised of a survey MS1 scan (mass range: 395-1005 m/z), with an accumulation time of 100 ms, a declustering potential of 80 V, and a collision energy of 10 V. MS2 scans were acquired using 80 variable width windows across the precursor ion mass range (250 -1500 m/z) , with an MS2 accumulation time of 25 ms, dynamic collision energy enabled, charge state 2 selected, and Zeno pulsing enabled (total cycle time 2.5 seconds).

### 2.6 | DIA data processing with Spectronaut

All data files were processed with Spectronaut v18 (version 18.6.231227.55695; Biognosys) performing a library-free search using the directDIA algorithm with the *Homo sapiens* Uniprot reference database with 20,526 entries, accessed on 08/28/2015. Dynamic data extraction parameters and precision iRT calibration with local non-linear regression were used. Trypsin/P was specified as the digestion enzyme, allowing for specific cleavages and up to two missed cleavages. Methionine oxidation, protein N-terminus acetylation, and deamidation of asparagine were set as dynamic modifications, while carbamidomethylation of cysteine was set as a static modification. Protein group identification (grouping for protein isomers) required at least 2 unique peptides and was performed using a 1% q-value cutoff for both the precursor ion and protein level. Protein quantification was based on the peak areas of extracted ion chromatograms (XICs) of 3 – 6 MS2 fragment ions, specifically b-and y-ions, with automatic normalization and 1% q-value data filtering applied. Relative protein abundance changes were compared using the Storey method with paired t-tests and p-values corrected for multiple testing using group wise testing corrections.(13)

## 3 | RESULTS AND DISCUSSION

### 3.1 | Proteomic workflow for young and old AML patients’ samples

The workflow in Figure 1 outlines the comprehensive approach taken to investigate age-related differences in AML cells between young and old patients. The known biological hallmarks of aging (4, 5), such as increased genomic instability, reduced mitochondrial quality, impaired autophagy, and heightened inflammatory responses, have been linked to age-associated vulnerabilities in AML progression (Figure 1A). The differential sensitivity to DNA-damaging agents and metabolic inhibitors observed between young and old HSCs emphasizes the importance of understanding these cellular aging mechanisms in AML pathogenesis. In our study design we obtained mononuclear cells from bone marrow aspirates or leukapheresis of old (n = 9) and young (n = 5) AML patients which were then subjected to a robust mass spectrometry workflow to quantify the proteomes (Figure 1B). In all samples the AML blasts made up the majority of cells present. The proteomic workflow (Figure 1C) was comprised of cell lysis, protein digestion, and microflow reverse-phase high performance liquid chromatography tandem mass spectrometry (LC-MS/MS) (9, 14, 15). Samples were analyzed using comprehensive data-independent acquisition (DIA) and data was processed using directDIA spectral library-free quantification. MS2 quantification provided insights into relative protein abundance changes for the 4,471 protein groups identified with at least two unique peptides (Table S1).

**Figure 1.**
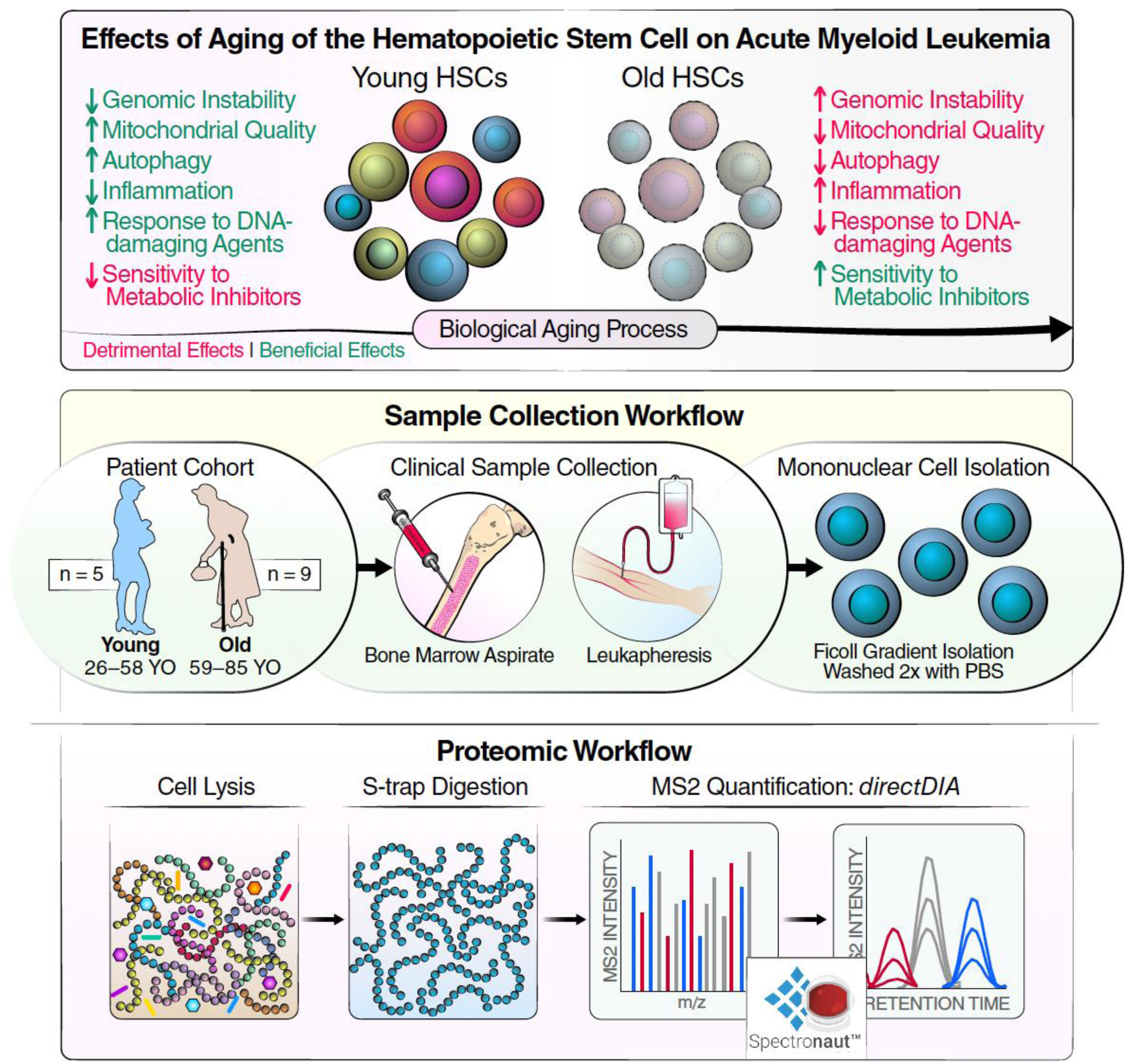
Workflow for investigating the effects of aging on hematopoietic stem cells (HSCs) in acute myeloid leukemia (AML) patients. (A) Schematic representation of the biological aging process in young and old HSCs, highlighting key factors such as genomic instability, mitochondrial quality, autophagy, inflammation, and response to DNA-damaging agents and metabolic inhibitors. (B) Sample collection workflow, detailing the collection of bone marrow aspirates and leukapheresis from young (n = 5, aged 26–58) and older (n = 9, aged 66–85) AML patients, followed by mononuclear cell isolation using Ficoll gradient centrifugation. (C) Proteomic workflow illustrating the S-Trap-based digestion, cell lysis, and MS2 quantification using directDIA for proteomic analysis of the isolated cells.

### 3.2 | Protein Profile Changes in AML Patients with Age (old vs young)

The primary focus of this work was to understand age-associated changes underlying AML. The partial least squares discriminant analysis (PLS-DA) plot (Figure 2A) illustrated the clear separation between old and young AML patient groups with a clear distinction of their proteomes. This separation suggested that age-associated changes in the proteome play a significant role in AML pathology. The proteomic comparison between old and young AML patients revealed distinct differences in protein abundance patterns. When comparing older and younger AML patient samples, 508 protein groups were significantly upregulated, and 381 protein groups were significantly downregulated (Figure 2B). Overall, the workflow featured an approach that, when combined with a diverse patient cohort, allowed for a thorough investigation of the proteomic changes associated with age in AML patients.

**Figure 2.**
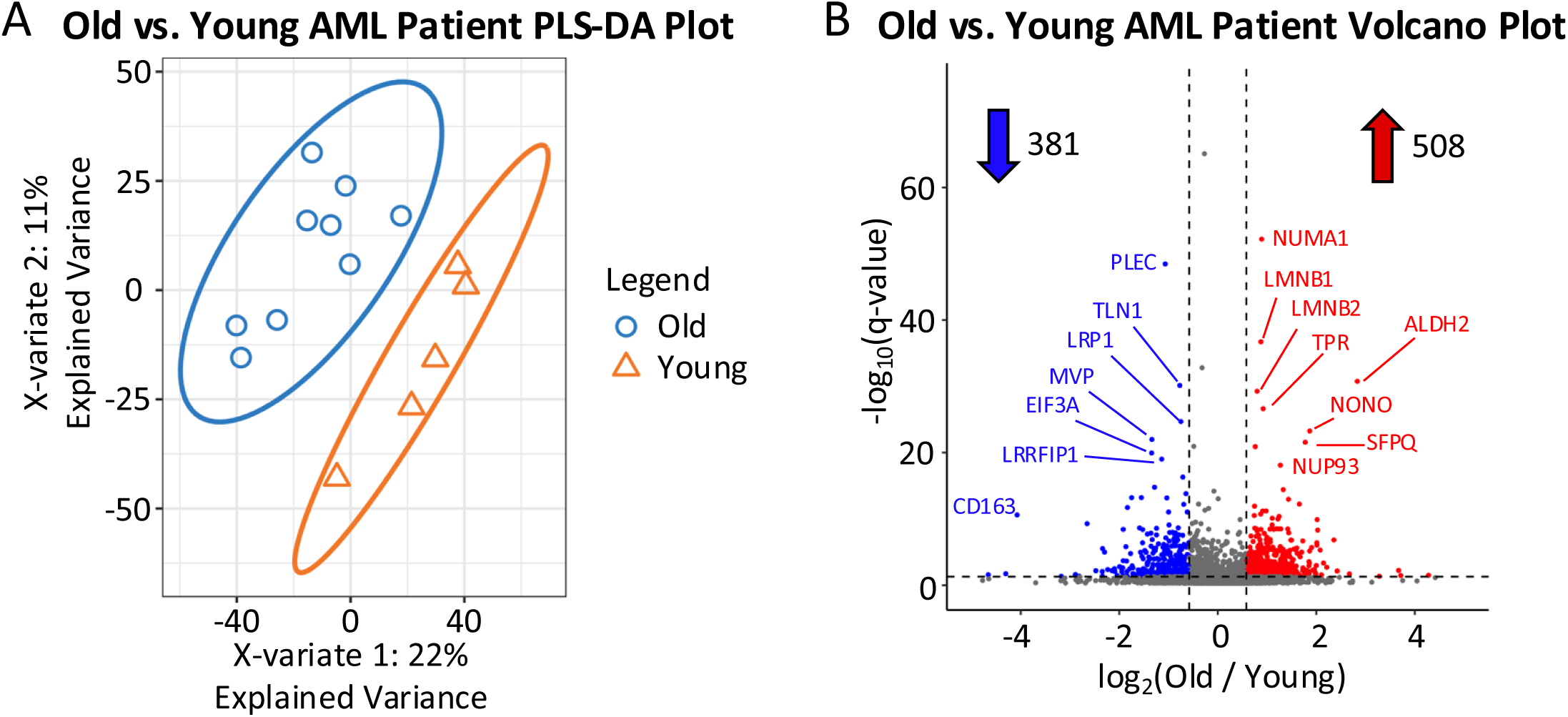
Comparative analysis of protein expression between old and young AML (acute myeloid leukemia) patients. (A) PLS-DA (Partial Least Squares Discriminant Analysis) plot depicting the separation of proteomic profiles between the two age groups based on X-variate 1 (22% explained variance) and X-variate 2 (11% explained variance). (B) Volcano plot showing differentially expressed proteins between old and young AML patients. Log2 fold changes (Old vs. Young) are plotted against the -log10 of the q-value (the plot y-axis is zoomed-in and the protein Neuroblast differentiation-associated protein AHNAK with q-value = 1.81e-116 is not displayed).

### 3.3 | Localization of Organellar Protein Changes and Gene Ontology Analysis

To highlight organelles that are most impacted by age in AML patients, we visualized the subcellular distribution of proteins by overlaying the fold change data from the old and young comparison on to a previously published map of 5,020 proteins assigned to 12 subcellular locations in human U-2 OS organelle lysates from a published localization of organellar proteins by isotope tagging (LOPIT) experiment using t-SNE coordinates (16, 17). A total of 3,192 proteins from the old and young comparison (investigated in this study) were overlayed onto the LOPIT plot (Figure 3A). One notable observation from the t-SNE plot was the pronounced upregulation of nuclear proteins in older AML patients compared to younger counterparts. This increase in nuclear protein abundance may be linked to age-related changes in nuclear architecture and chromatin remodeling. Mitochondrial proteins also showed distinct upregulation in older patients, which potentially reflects age-associated mitochondrial dysfunction and altered energy metabolism, a hallmark of aging and age-related diseases like AML. In contrast, proteins associated with the lysosome, endoplasmic reticulum, and plasma membrane showed more subtle abundance differences, indicating that aging may have a lesser impact on these compartments in AML. Older AML patients showed a distinct downregulation of cytosolic, ribosome 40S, and ribosome 60S proteins when comparing to young AML patients (Table S2). This may suggest that while cellular processes like translation and protein metabolism (which are handled by ribosomes) are downregulated in older patients, the overall protein synthesis machinery in these cells may be compromised. The observed decrease in ribosomal proteins points to reduced translational capacity, which is a key feature of aging cells.

**Figure 3.**
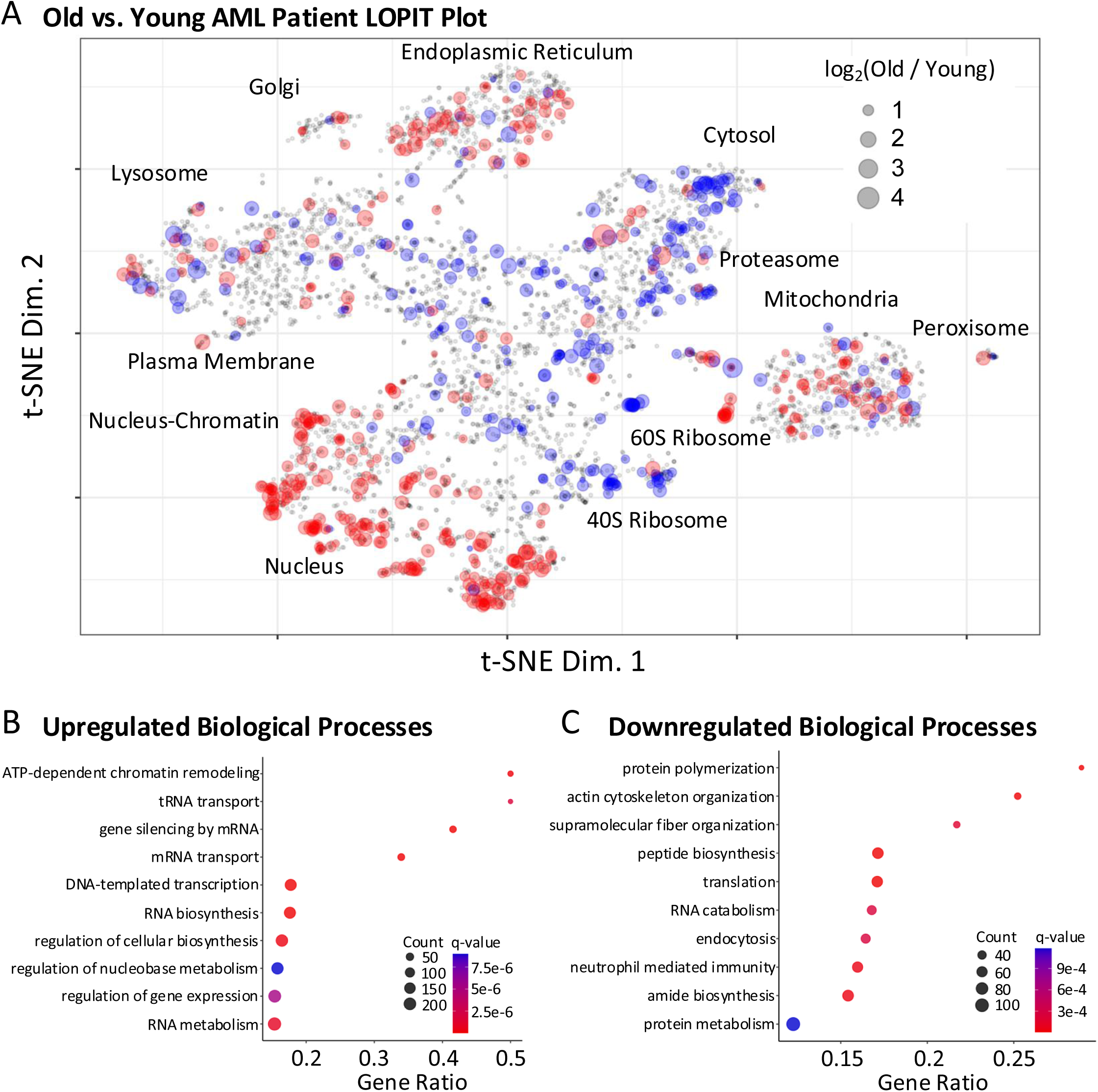
Localization of organelle proteomes in old versus young AML patients using LOPIT (Localization of Organelle Proteins by Isotope Tagging). (A) t-SNE plot representing the spatial distribution of proteins across different subcellular compartments between old and young AML patient samples. Organelles analyzed include the nucleus, mitochondria, lysosome, and endoplasmic reticulum. (B) Upregulated biological processes in older AML patients include ATP-dependent chromatin remodeling, mRNA transport, and gene silencing by RNA. (C) Downregulated processes include actin cytoskeleton organization, translation, and protein metabolism. Gene ratio, q-value, and count are plotted for both upregulated and downregulated biological processes.

The analysis of biological processes, using ConsensusPathDB (18) (Figure 3B-C), revealed that ATP-dependent chromatin remodeling, mRNA transport, and gene silencing by RNA were significantly upregulated in older patients, pointing to enhanced transcriptional and post-transcriptional regulatory mechanisms. The upregulated regulatory processes are crucial for maintaining cellular homeostasis and genomic stability, which may become dysregulated in the aging AML microenvironment. On the other hand, processes such as actin cytoskeleton organization, translation, and protein metabolism were markedly downregulated, suggesting cytoskeletal dynamics and protein synthesis impairment in the older cohort. These findings underscored the complexity of age-related proteomic alterations in AML and highlighted key pathways that may serve as potential therapeutic targets for addressing age-associated vulnerabilities in AML. The localization shifts in mitochondrial, nuclear, and cytoskeletal proteins further suggested that cellular aging in AML may involve a coordinated disruption of both energy metabolism and structural integrity, contributing to the pathophysiology of the disease in older patients.

A decline in protein synthesis may lead to diminished cellular resilience, reduced capacity for damage repair, and impaired ability to produce proteins critical for cellular homeostasis. This downregulation may also reflect a broader reorganization of metabolic priorities in older AML patients, where cells might favor energy conservation over protein production due to the impaired function of metabolic pathways such as mitochondrial respiration. Furthermore, diminished ribosomal activity could contribute to reduced protein turnover, exacerbating the accumulation of damaged or misfolded proteins, which is commonly seen in aged cells and may enhance the leukemic phenotype in older patients. These findings are consistent with the broader concept of translational attenuation in aging, which has been linked to cellular senescence and impaired stress responses. Understanding the specific molecular consequences of ribosomal downregulation could guide the development of age-targeted therapeutic strategies that aim to restore or compensate for impaired translational processes in older AML patients.

### 3.4 | Upregulation of RNA Splicing Factors in Older AML Patients

Our proteomic analysis identified major splicing regulators, including core spliceosomal components, RNA-binding proteins and heterogeneous nuclear ribonucleoproteins, RNA helicases, transcription-coupled splicing factors, THO/TREX complex members, and cleavage/polyadenylation factors, highlighting the complex interplay between various aspects of RNA processing machinery in AML with age. Of the identified 69 proteins that were related to mRNA splicing or the spliceosome, a large proportion of proteins in the splicing machinery were significantly upregulated in older AML patients compared to younger patients (Figure 4; Table S3). Among the significantly altered proteins, non-POU domain-containing octamer-binding protein (NONO, FC = 3.65, q = 5.47E-24) and splicing factor proline and glutamine rich (SFPQ, FC = 3.42, q = 2.70E-22) were the most upregulated splicing factors in older AML patients compared to their younger counterparts. Upregulation of NONO has been shown to reduce the sensitivity of AML cells to the chemotherapy agent Cytarabine. (19) Protein SFPQ is involved in multiple functions in processing of pre-mRNA and transcriptional regulation with histone deacetylases which may play a role in deregulation of splicing in leukemic cells.(20)

**Figure 4.**
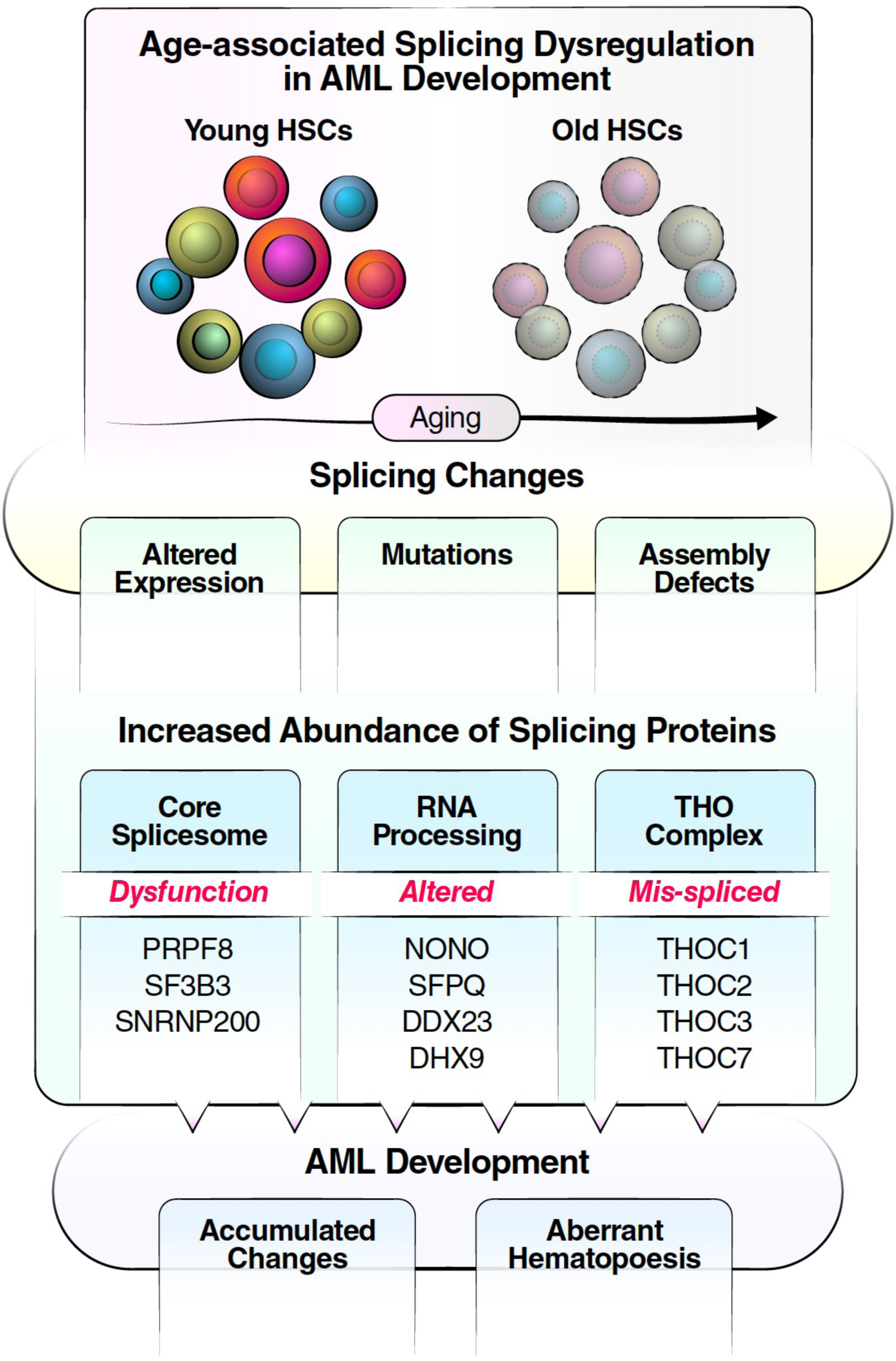
Age-associated dysregulation of splicing in AML. Proteomic analysis of hematopoietic stem cells from patients with acute myeloid leukemia (AML) revealed a dramatic alteration of the splicing machinery in older patients. Patient Age was associated with dysregulation in the splicing machinery, that featured coordinated up-regulation of core spliceosome, RNA processing-associated proteins, and THO complex proteins, which may contribute to specific or more detrimental AML pathogenesis in older patients.

Additionally, ubiquitin carboxyl-terminal hydrolase 39 (USP39, FC = 4.23, q = 5.06E-06) and pre-mRNA-processing factor 40 homolog A (PRPF40A, FC = 3.26, q = 1.02E-05) also showed substantial upregulation, suggesting enhanced splicing activity in older AML patients. These proteins are known components of the spliceosome, a critical complex involved in the removal of introns from pre-mRNA. USP39 abundance was previously correlated with poor survival of patients with leukemia.(21) The elevated abundance of these splicing factors suggests widespread changes in RNA processing, particularly in alternative splicing mechanisms. Such dysregulation may lead to the generation of oncogenic splice variants or the aberrant splicing of tumor suppressor genes, thereby contributing to leukemogenesis in older patients.(22) Conversely, several splicing factors were significantly downregulated in the older AML patient cohort (vs younger AML patients). Notably, neuroblast differentiation-associated protein (AHNAK, FC = 0.33, q = 1.81E-116) exhibited the most pronounced downregulation, followed by poly(A) binding protein cytoplasmic 1 (PABPC1, FC = 0.56, q = 8.66E-05) and aquarius intron-binding spliceosomal factor (AQR, FC = 0.50, q = 4.62E-03). AHNAK has been implicated in the stimulation of adipocyte differentiation and has previously been found to be under-expressed in AML patients.(23) PABPC1 plays a crucial role in cytoplasmic mRNA homing and regulates almost every aspect of RNA metabolism.(24) The diminished expression of these proteins may reflect age-related declines in cellular homeostasis, contributing to the altered splicing landscape observed in older AML patients. Supporting these findings are reports of frequent overexpression of splicing factor genes, widespread splicing alterations, and splicing factors associated with RNA metabolism in AML patients compared with healthy controls.(25–27) Most importantly, our study demonstrated that overexpression of proteins associated with splicing is even higher in older AML patients compared to younger AML patients highlighting the key relevance of aging during AML disease progression.

### 3.5 | Dysregulation of Splicing Machinery in Older AML Patients

Multiple components of the RNA polymerase II complex (POLR2A, POLR2B, POLR2C, POLR2D, POLR2E, POLR2G, and POLR2H) were also found to be upregulated in older AML patients compared to young AML patients. These polymerase subunits are essential for transcription and co-transcriptional splicing, suggesting that increased transcriptional activity is coupled with aberrant splicing processes in older AML patients. The concurrent upregulation of transcriptional regulators, such as RNA binding protein fused in sarcoma (FUS) and SON DNA and RNA binding protein (SON), which are involved in chromatin remodeling and transcriptional elongation, points to an epigenetic dimension in the dysregulated gene expression observed in this cohort. FUS-transcription factor ERG (ERG), a chimeric gene associated with poor outcomes in myelodysplastic syndromes and acute myeloid leukemia, induces resistance to the chemotherapy drug azacitidine by disrupting chromatin organization through co-repressors.(28) SON regulates transcription initiation, but upregulated short isoforms in AML disrupt this repression and serve as markers of abnormal transcriptional initiation.(29) The integration of transcriptional and splicing dysregulation likely contributes to the complex gene expression profiles that drive leukemogenesis and disease progression in older AML patients.

Splicing-associated proteins involved in RNA stability and transport, including six members of the heterogeneous nuclear ribonucleoprotein family (HNRNPC, HNRNPH1, HNRNPH3, HNRNPK, HNRNPR, and HNRNPUL1) and two subunits of the cleavage stimulation factor (CSTF2 and CSTF3), exhibited significant increase and upregulation in older AML patients (fold changes ranging from 1.63 to 3.50). These proteins are key regulators of mRNA processing and stability, ensuring that mRNAs are properly exported from the nucleus and remain stable in the cytoplasm for translation. The increased abundance of these proteins, previously demonstrated to regulate cellular proliferation,(30, 31) suggests that AML cells in older patients may rely on enhanced RNA stability and export to sustain high levels of oncogenic proteins. This reliance likely supports sustained leukemia cell proliferation and survival, contributing to the more aggressive disease phenotype often observed in older AML patients. The increased abundance of these RNA processing proteins may provide potential therapeutic targets, as disrupting their function could impair the stability and translation of oncogenic mRNAs, thereby inhibiting leukemia progression.

### 3.6 | Chromatin Remodeling and Epigenetic Regulation

In addition to transcriptional and splicing dysregulation, our study identified significant upregulation of chromatin-remodeling proteins, such as A-kinase anchoring protein 8 like (AKAP8L) and THO complex (THOC) proteins in older AML patients. AKAP8L is known to regulate mitosis and cell growth,(32) and its increased expression may facilitate the unchecked proliferation characteristic of leukemic cells. The concurrent upregulation of THOC proteins (THOC1, THOC2, THOC3, and THOC7) highlights a critical link between transcriptional regulation and post-transcriptional RNA processing in older AML patients (Figure 4). THOC proteins play a pivotal role in mRNA export, telomere maintenance, and chromatin organization. The increased expression of these proteins suggests a coordinated effort to sustain altered gene expression profiles that favor leukemogenesis. Specifically, THOC-mediated regulation of telomeres and chromatin structure may lead to genomic instability and persistent epigenetic changes, facilitating the survival and expansion of leukemic clones.(33) The upregulation of chromatin-remodeling proteins in older AML patients indicates that epigenetic modifications are likely driving aberrant gene expression patterns. Persistent changes in chromatin structure can result in the activation of oncogenes and repression of tumor suppressor genes, thereby contributing to disease progression and resistance to conventional therapies. This epigenetic landscape may also interact with transcriptional and splicing dysregulation, creating a complex network of regulatory disruptions that promote leukemogenesis.

## 4 | CONCLUDING REMARKS

This comprehensive proteomic analysis uncovered significant age-related molecular alterations in hematopoietic stem cells (HSCs) of AML patients, highlighting the complex interplay between aging and leukemia pathogenesis. Older AML patients (compared to younger AML patients) exhibited a distinct proteomic profile marked by the upregulation of nuclear and mitochondrial proteins, indicative of increased genomic instability and mitochondrial dysfunction—hallmarks of cellular aging. Simultaneously, the downregulation of cytosolic and ribosomal proteins suggested a potential decline in protein synthesis and translational capacity, potentially reducing cellular resilience and impairing homeostasis in aged leukemic cells. Gene ontology and localization analyses revealed coordinated disruptions in critical cellular processes, including energy metabolism, transcriptional regulation, and structural integrity within the aging AML microenvironment. The heightened expression of chromatin-remodeling and epigenetic regulators in the aged AML cohort may drive some of the aggressive disease phenotypes often observed in older patients. Additionally, aberrant RNA splicing patterns, independent of splicing factor mutations, are strongly linked to poor prognosis by activating oncogenic pathways and fostering treatment resistance.

A key finding in the older AML cohort was the upregulation of splicing factors - and overall dysregulation of the complex splicing machinery - alongside dysregulated transcription, enhanced RNA stability, and activated DNA repair pathways in older AML patients (Figure 4). These molecular changes contributed to the unique biology of AML in the older, underpinning increased chemoresistance and poor prognosis. Specifically, elevated splicing factors such as NONO, SFPQ, USP39, and PRPF40A, along with enhanced transcriptional machinery, facilitate the generation of oncogenic splice variants and aberrant gene expression, promoting leukemogenesis and disease progression. The involvement of survival-associated alternative splicing in essential cellular functions, such as protein translation and stress response, underscores their role in treatment resistance and disease aggressiveness. Targeting these splicing mechanisms presents a promising therapeutic strategy to overcome chemoresistance and improve clinical outcomes for older AML patients. By disrupting dysregulated pathways—such as splicing factors, transcriptional regulators, and chromatin remodelers—age-tailored interventions can restore normal cellular functions and enhance treatment efficacy.

Our study elucidated the intricate landscape of RNA splicing and transcriptional dysregulation in older AML patients, revealing multifaceted strategies employed by leukemic cells to sustain growth and resist treatment. These molecular insights not only enhanced our understanding of AML pathogenesis in older individuals but also identify potential prognostic markers and therapeutic targets. Future research may focus on assessing these findings also in larger human cohorts and developing targeted interventions to disrupt these dysregulated pathways, ultimately improving outcomes for older AML patients. In summary, this study provided a detailed molecular landscape of age-related changes in AML, emphasizing the complex regulatory networks that drive disease progression in older individuals. By identifying key proteomic shifts and their associated biological processes, we pave the way for the development of targeted, age-tailored therapies that offer more effective and personalized treatment approaches, thereby potentially improving clinical outcomes for older AML patients.

## Funding information

Pilot funds of the Comprehensive Cancer Center of Atrium Health Wake Forest Baptist.

## ACKNOWLEDGEMENTS

We are grateful for the generous support for the ZenoTOF 7600 platform from SCIEX.

## CONFLICTS OF INTEREST

Birgit Schilling is serving on the Advisory Board for MOBILion Systems. Timothy Pardee is a member on the following Advisory Boards: AbbVie, Genentech, AstraZeneca, Ipsen, and he received research support from Research Support from Karyopharm, Delta Fly Pharmaceuticals, and NCI. The other authors declare no conflicts of interest.

## DATA AVAILABILITY STATEMENT

The raw data and complete mass spectrometry data sets have been uploaded to the Mass Spectrometry Interactive Virtual Environment (MassIVE) repository, which is maintained by the Center for Computational Mass Spectrometry at the University of California San Diego. These data can be accessed and downloaded using the following ftp link: ftp://MSV000097444@massive.ucsd.edu or via the MassIVE website: https://massive.ucsd.edu/ProteoSAFe/dataset.jsp?task=1d6fa995249e43d196e86c001d <u>73c31a</u> (MassIVE ID number: MSV000097444; ProteomeXchange ID: PXD062218). [Note to the reviewers: To access the data repository MassIVE (UCSD) for MS data, please use: Username: MSV000097444_reviewer; Password: winter].

## Abbreviations

AML: acute myeloid leukemia
ATP: adenosine triphosphate
CAD: collision associated dissociation
DIA: data-independent acquisition
DMSO: dimethyl sulfoxide
DNA: deoxyribonucleic acid
DTT: dithiothreitol
EDTA: ethylenediaminetetraacetic acid
FBS: fetal bovine serum
FC: fold change
HLB: hydrophilic–lipophilic balance
HSC: hematopoietic stem cells
IAA: iodoacetamide
iRT: indexed retention time
LC-MS/MS: liquid chromatography tandem mass spectrometry
LOPIT: localization of organellar proteins by isotope tagging
mRNA: messenger ribonucleic acid
PIC: protease/phosphatase inhibitor cocktail
PLS-DA: partial least squares discriminant analysis
SDS: sodium dodecyl sulfate
TEAB: triethylammonium bicarbonate buffer
WBC: white blood cell
XIC: extracted ion chromatogram

